# Temperature-dependent changes in neuronal dynamics in a patient with an *SCN1A* mutation and hyperthermia induced seizures

**DOI:** 10.1101/048520

**Authors:** C Peters, RE Rosch, E Hughes, P Ruben

**Affiliations:** Department of Biomedical Physiology and Kinesiology, Simon Fraser University, Burnaby, BC, Canada; Wellcome Trust Centre for Neuroimaging, Institute of Neurology, University College London, UK; Centre for Developmental Cognitive Neuroscience, Institute of Child Health, University College London, UK; Department of Paediatric Neurology, Evelina London Children’s Hospital, Guy’s and St Thomas’ NHS Foundation Trust, London, UK

## Abstract

Dravet syndrome is the prototype of *SCN1A*-mutation associated epilepsies. It is characterised by prolonged seizures, typically provoked by fever. We describe the evaluation of an *SCN1A* mutation in a child with early-onset temperature-sensitive seizures.

The patient carries a heterozygous miss-sense variant (c3818C>T; pAla1273Val) in the NaV1.1 brain sodium channel. We characterised the functional effects of the variant using patch clamp recordings on Chinese Hamster Ovary Cells at different temperatures (32, 37, and 40°C).

The variant channels produced a temperature-dependent destabilization of channel activation and fast inactivation. Implementing these empirical abnormalities in a computational model produces a higher threshold for depolarization block in the variant, particularly at 40°C, suggesting a failure to autoregulate at high-input states.

These results reveal direct effects of abnormalities in biophysical NaV1.1 channel properties on neuronal dynamics. They illustrate the value of combining cellular measurements with computational models to integrate different scales (gene/channel to patient) of observations.

## Introduction

Epilepsy is a common neurological condition, with particularly high incidence in early childhood ^1^. The diagnosis is made when patients show continued susceptibility for recurrent epileptic seizures. Although in some patients this susceptibility is due to structural brain abnormalities, including tumours, previous strokes, or congenital malformations, for others the suspected cause is genetic ^2^.

Mutations in a wide variety of ion channel and synaptic genes cause epilepsy in individual patients and families ^3^. Somewhat surprisingly, many of these mutations were identified not from common, familial epilepsies (recently renamed *genetic generalised epilepsies*) ^4^, but from epileptic encephalopathies ^5^, sporadic severe epilepsies occurring mainly in childhood.

One of the first ion channel “epilepsy” genes identified was *SCN1A*, which encodes the neuronal voltage-gated sodium channel NaV1.1. Initially described in patients with Generalised Epilepsy with Febrile Seizures plus Type 2 (GEFS+ Type 2) ^6^, it is also the dominant genetic cause of Dravet syndrome, previously termed severe myoclonic epilepsy of infancy (SMEI) ^7^, an early onset epileptic encephalopathy. The link between the two conditions was made because of the shared susceptibility to recurrent, prolonged febrile seizures; Dravet patients in particular experience seizures in response to increases in environmental temperatures, such as hot baths ^8^.

*SCN1A* mutations related to Dravet syndrome include severe disruptions of channel integrity (e.g. frameshift mutations, deletions), and, less commonly, missense mutations leading to either channel impairment or gain of function. The reported prevalence of loss-of-function mutations in clinical cohorts ^9^ may be counterintuitive, since they would render neurons less excitable and thus less seizure-prone. This paradox may be explained by studies suggesting NaV1.1 channels are predominantly found in GABAergic interneurons, where loss-of-function may cause overall disinhibition, permissive of epileptic activity ^10,11^. This also explains the paradoxical exacerbation of seizures in response to sodium channel blocking antiepileptic drugs observed in many Dravet patients ^12^.

This NaV1.1 haploinsufficiency account of epileptogenesis in Dravet and associated epilepsies does not fully explain all clinical observations. A significant number of patients with gain-of-function mutations show severe epilepsy phenotypes considered more typical of deletion or frameshift mutations ^9,13,14^.

Mapping possible direct mechanistic links between sodium channel mutations and increased seizure susceptibility may help improve our understanding of genotype-phenotype correlations. By integrating experimental measurements into a computational model of neuronal function, we can predict the effects of mutations on neurons *in silico*. This may identify otherwise unpredictable functional effects of genetic mutations and group abnormalities in the system into functionally relevant categories ^15,16^.

We present a patient with an early-onset, temperature-sensitive epilepsy phenotype and a *de novo* heterozygous *SCN1A* mutation (c.3818C>T, ClinVar Accession: RCV000180969.1) coding for a mutant in DIIIS2 of the NaV1.1 channel (p.Ala1273Val). Using patch-clamp characterisation of channel properties, we identify dynamic, temperature-dependent differences from wild type (WT). Integrating these empirical results in computational models of action potential dynamics at the neuronal membrane, we specify the functional effects of the mutation and describe a mechanism that leads to temperature-sensitive epilepsy.

## Results and Discussion

Sample macroscopic sodium currents from WT and A1273V channels are shown in figures 1A and 1B, respectively. There is no significant difference in the time to 50% maximal current between WT and A1273V channels nor is there a difference in temperature sensitivity (Figure 1C and 1D). Increasing temperature significantly accelerates the time to 50% maximal current at potentials between −20 mV and +50 mV (P = 0.0006 to 0.0215) in both WT and A1273V. There is a significant difference in temperature sensitivity of the conductance-voltage relationship between WT and A1273V (P = 0.0235). When temperature is increased from 37 °C (Figure 2A) to 40 °C (Figure 2B), the A1273V activation curve midpoint is depolarized by 8 mV whereas the WT is depolarized by only 4.5 mV.

**Figure 1:**
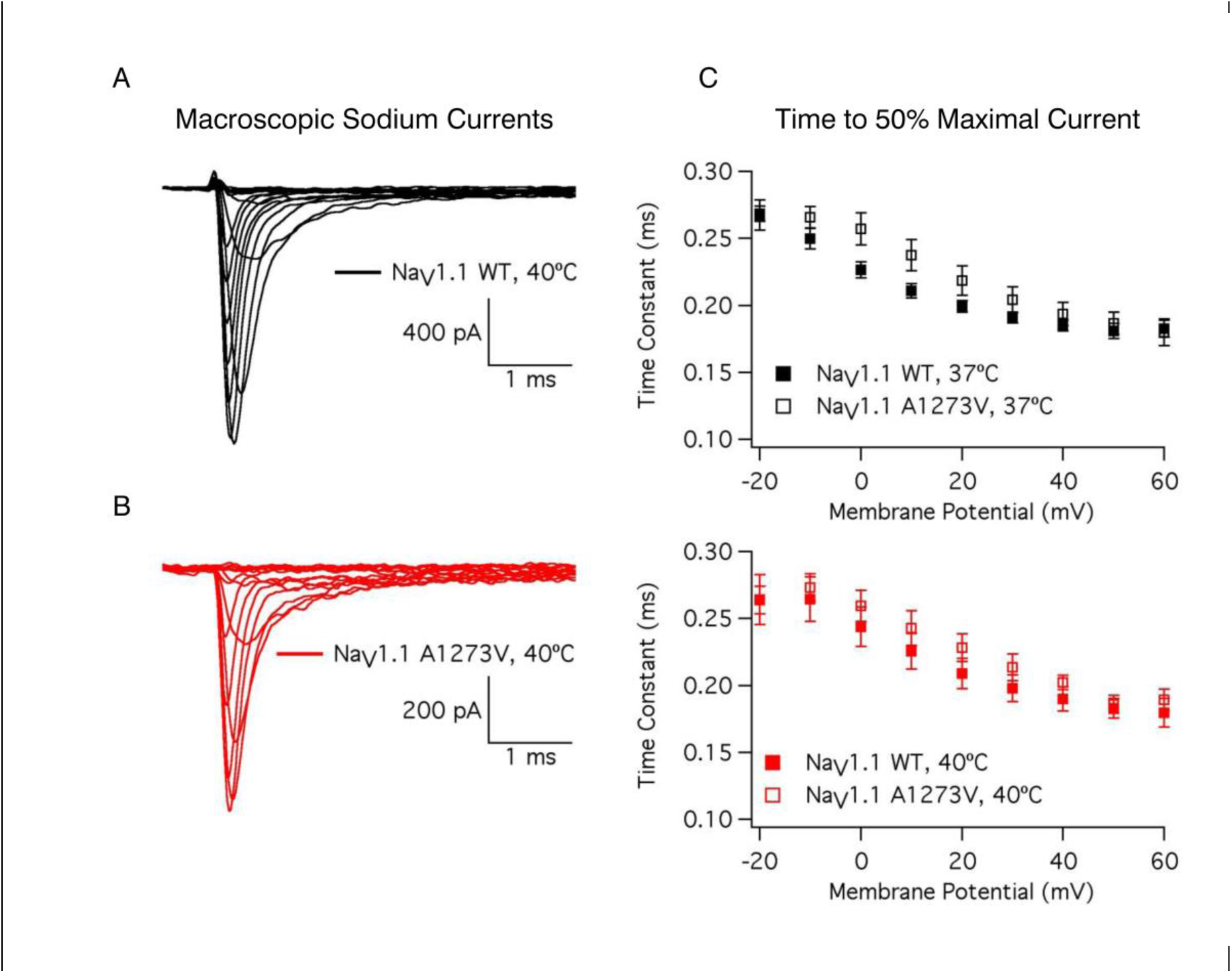
Macroscopic NaV1.1 Currents. Sample WT (A) and A1273V (B) currents elicited at potentials between −100 mV and +60 mV at 40 °C. Time to 50% maximal current is plotted versus voltage for WT and A1273V NaV1.1 channels at 37 °C (C) and 40 °C (D).

**Figure 2:**
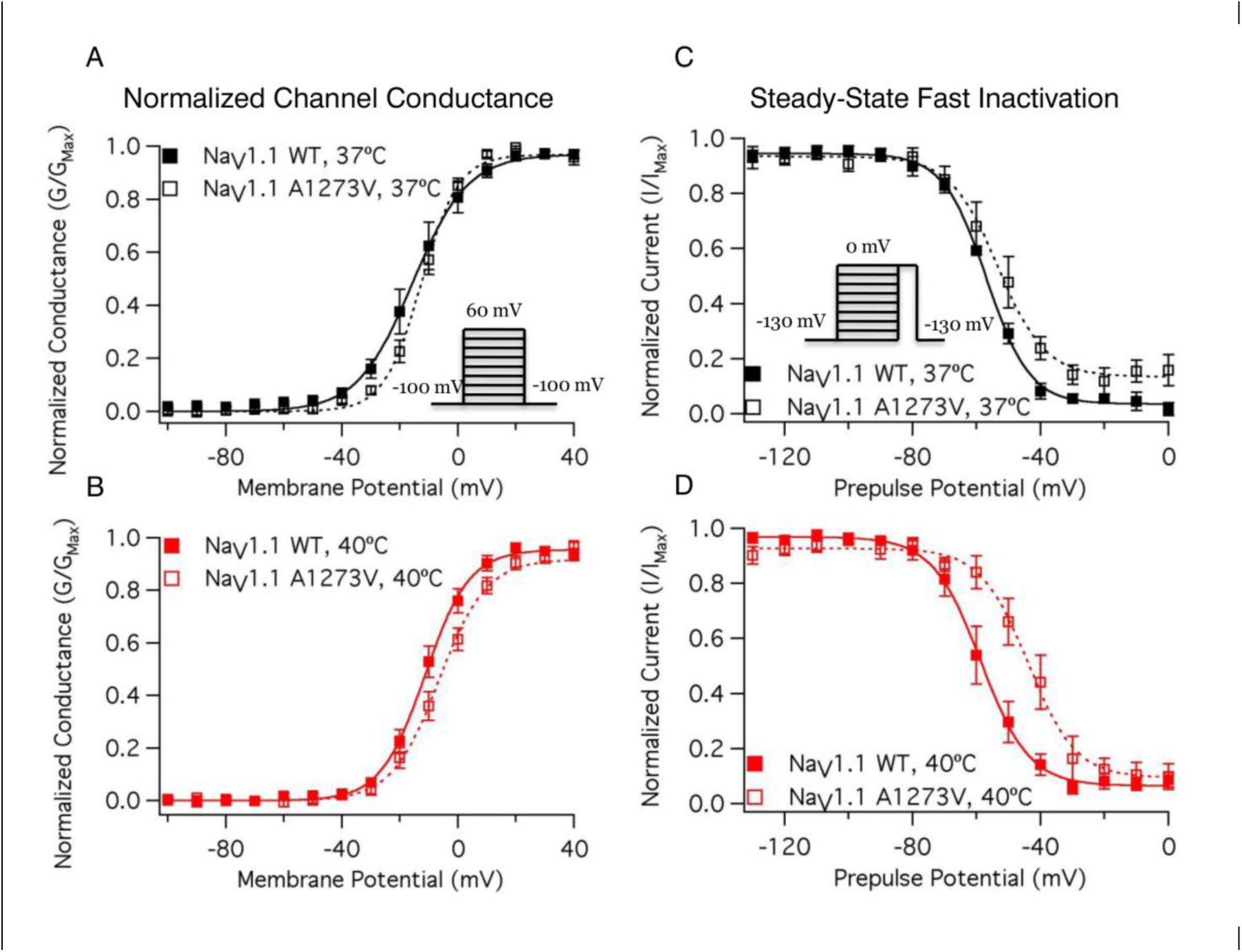
Voltage Dependence of NaV1.1 Conductance and Fast Inactivation. Normalized Conductance curves for WT and A1273V NaV1.1 channels at 37 °C (A) and 40 °C (B). Conductance was determined from macroscopic current recordings using Ohm’s law corrected for the experimentally observed equilibrium potential. Normalized current during a test pulse following a 200 ms pre-pulse is plotted versus pre-pulse potential for WT and A1273V NaV1.1 channels at 37 °C (C) and 40 °C (D). Pulse protocols used to elicit macroscopic current and to determine the voltage-dependence of fast inactivation are shown in the insets of A and C, respectively.

There is a significant difference in the temperature sensitivity of the midpoint of steady-state fast inactivation between WT and A1273V (P < 0.0001). At 37 °C the A1273V steady-state fast inactivation midpoint is depolarized by 1 mV compared to WT (Figure 2C); the difference increases to 14 mV at 40 °C (Figure 2D). The fast time constant of recovery is significantly accelerated by increases in temperature (P = 0.0028); the slow time constant of recovery is not (P = 0.8631). Neither the fast nor slow time constants of recovery are altered by the mutation. There is a significant difference in the temperature sensitivity of the recovery component amplitudes in the A1273V mutant compared to WT (P = 0.0021). Increasing temperature decreases the fast component amplitude and increases the slow component amplitude in WT channels; the opposite occurs in A1273V. The overall results are an increased recovery in A1273V (Fig. 3B) channels at 40 °C compared to WT (Fig. 3A) channels. Open state fast inactivation time constants are shown for WT and A1273V channels in figures 3C and 3D, respectively. A mutant effect on fast inactivation onset occurs only at 0 mV (P = 0.0366); we conclude that A1273V has minimal impacts on fast inactivation onset.

**Figure 3:**
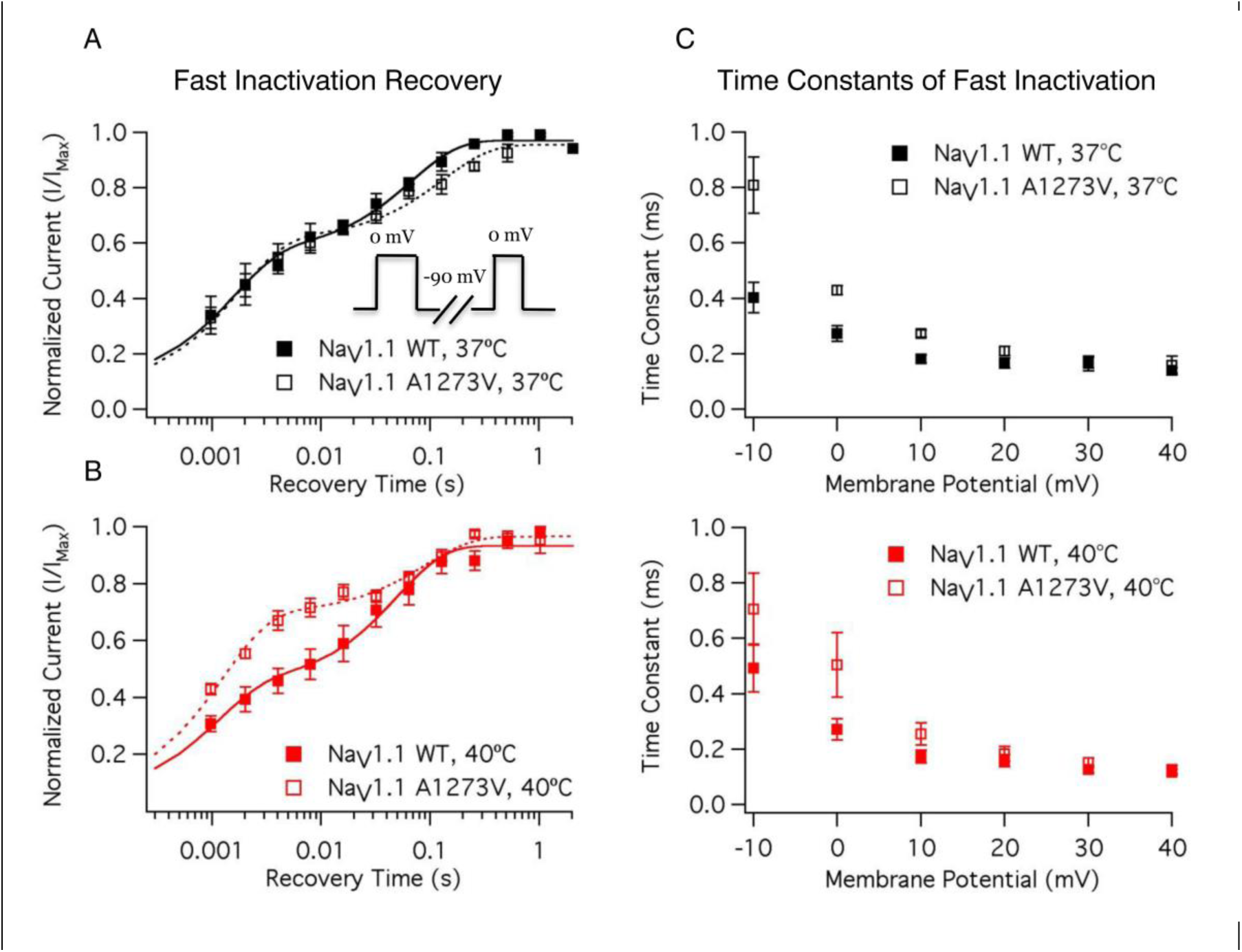
Time Course of Fast Inactivation. Open-state fast inactivation time constants are plotted versus voltage for WT and A1273V NaV1.1 at 37 °C (A) and 40 °C (B). The time course of fast inactivation recovery versus recovery is plotted for WT and A1273V NaV1.1 at 37 °C (C) and 40 °C (D). The double-pulse protocol used to measure fast inactivation recovery is shown in the inset of A.

We implemented cortical neuron models incorporating WT and A1273V data at 37 and 40 °C. Simulations with minimal input (*I*_*stim*_ = 0.2, Eq. 1.1) reveal little difference in firing frequency or amplitude for either temperature condition (Fig 4A). As the input stimulus increases, the action potential firing rates of WT models increases faster than those of A1273V models (Supplementary Figure 1). The lowest firing rate at a given stimulus is that of the A1273V model at 40 °C, consistent with an increased threshold for action potential firing, predicted by the depolarized conductance-voltage relationship, which is a loss of function in the mutant channel.

**Figure 4:**
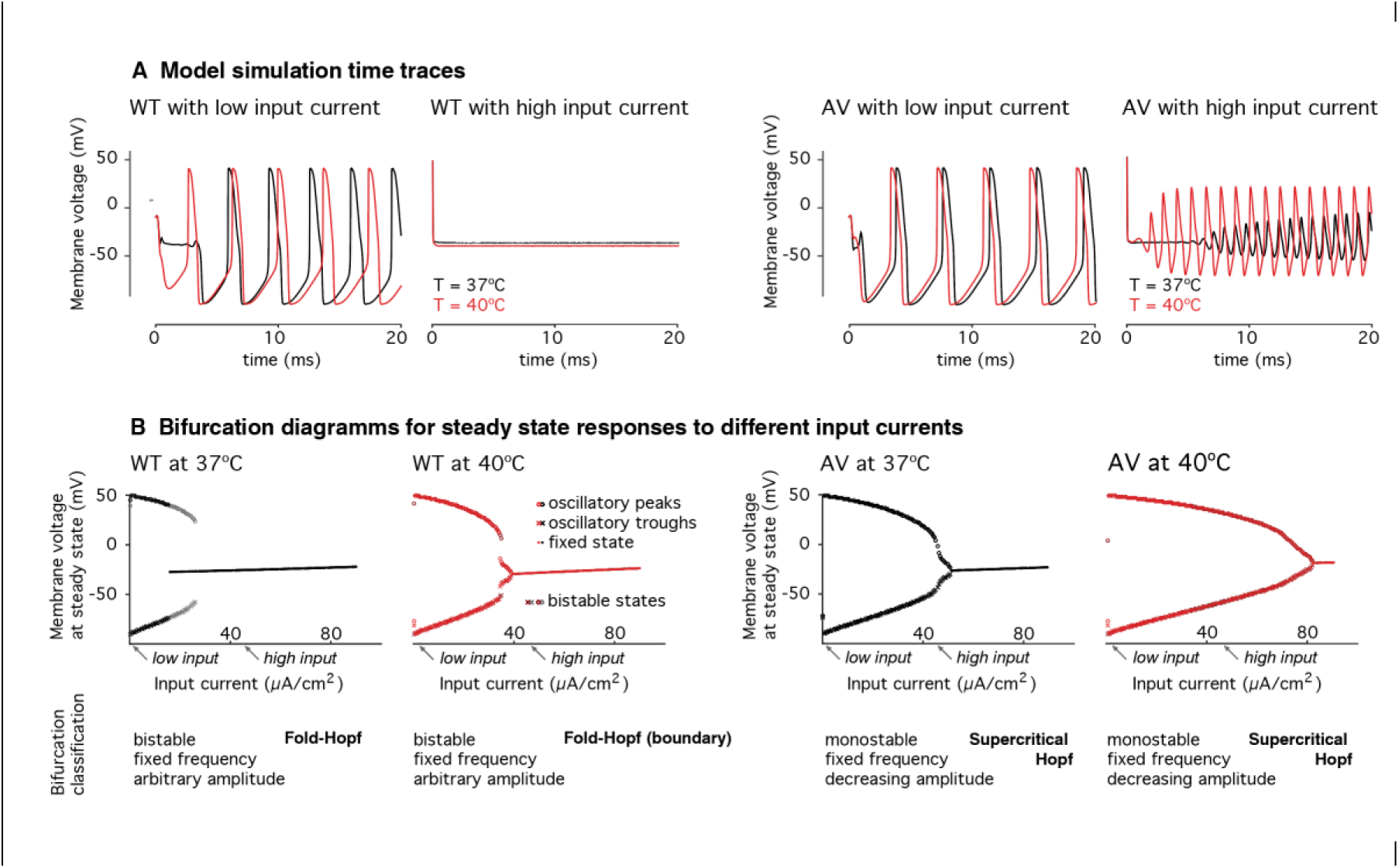
Computational modelling of membrane dynamics. Experimental voltage clamp measurements for four experimental conditions (WT and A1273V at 37°C and 40°C) were integrated into a Hodgkin-Huxley model of cortical neurons and normalised to the WT measurements at 37°C. (A) Simulations of the membrane response at different input current levels revealed absent action potential generation in the WT at high currents, even when A1273V neurons continue to fire. (B) Bifurcation analysis shows differences in the transition from oscillation to fixed steady states at high input currents (i.e. depolarisation block), both qualitatively in terms of the bifurcation type, and quantitatively in terms of the input currents required to achieve depolarisation block. Of all experimental conditions modelled, A1273V neurons permit the highest input currents to elicit continuous action potentials. **Model parameters**: *g*_*L*_ = *2.5*\**10*^−*5*^*S/cm*^*2*^, *E*_*L*_ = -*70.3*, *g*_*Na*_ = *0.056 S/cm*^*2*^, *E*_*Na*_ = *50* -*mV*, *g*_*K*_ = *0.005 S/cm*^*2*^, *E*_*K*_ = -*90 mV*, *V*_*t*_ = -*60 mV*, *C*_*m*_ = *0.01 μF/mm*^*2*^; (^17^); high input current *I*_*stim*_ = *45μA/mm*^*2*^, low input current *I*_*stim*_ = *0.2 μA/mm*^*2*^. Remaining parameters were condition specific and defined as described in the Methods section.

Simulations with high input currents (*I*_*stim*_ = *45μA/mm*^*2*^, Eq. 1.1), reveal significant divergence in the dynamic behaviours of different model neurons. Although there is no response from the WT neurons at either temperature (due to depolarisation block), there is continued action potential firing in A1273V neurons at both temperatures, with higher amplitude firing at 40°C. Depolarisation block describes the observation that neurons stop responding at high levels of stimulation despite being depolarised beyond the firing threshold. There is evidence that it might play a role in seizure termination ^18^,^19^.

Transitions between fixed steady states and oscillations (birfurcations) characterise the dynamic behaviour of neuronal oscillators ^20^. Comparing A1273V to WT, there is a shift of the oscillation offset bifurcation towards higher input current values, particularly at 40°C (Figure 4B). This means A1273V neurons in the hyperthermic condition continue to produce action potentials at high input currents. These data may best be explained by the depolarizing shift in the steady-state fast inactivation curve leading to increased channel availability in A1273V at 40 °C at a given membrane potential.

This study highlights the value of characterising *SCN1A* variants at elevated temperatures. One previous study identified a single-temperature sensitive loss-of-function mutant, R865G, in the NaV1.1 channel ^21^. Febrile seizures have also been successfully elicited in mouse models of Dravet syndrome ^22^. Our study suggests that a single mutation can confer both temperature-sensitive loss-of-function (i.e. lower action potential frequency) and gain-of-function (i.e. action potential generation at higher input currents) in Dravet syndrome. This approach adds a mechanistic understanding to existing evidence that even single, stem cell derived neurons carrying loss-of-function mutations show a hyperexcitable phenotype ^23^.

Our findings show functional impairments of A1273V are temperature-specific, suggesting a link between the phenotype and the genetic mutation. Embedding these functional abnormalities at the channel level into a computational neuronal model suggests that, while low input stimulation leads to decreased action potential firing, the maximum stimulus current which results in oscillations is higher in the mutant at 40 °C. Our model also suggested some more subtle differences. In the WT, there are bistable bifurcations in which neurons that enter depolarisation block, are subsequently more likely to remain at a fixed (i.e. blocked) steady state when stimulating currents decrease. In models incorporating the mutant channel dynamics, bifurcations are monostable; neurons which enter depolarisation block return to oscillatory behaviour when stimulation falls even just below the bifurcation threshold. These findings suggest a pathophysiological mechanism in which neurons recover their excitability more quickly after excessive stimulation.

Combining empirical measurements of impairments in molecular function with computational neuronal modelling integrates different scales of evidence. A similar approach identified novel pathological mechanisms resulting in abnormally increased persistent sodium currents through mutant channels in *SCN1A* mutations ^24^. Computational models have helped identify a common mechanism of epilepsy pathophysiology: apparently functionally disparate mutations affecting different sodium channels can result in neuronal hyperexcitability ^25^.

Our study design allows for comprehensive assessment of the molecular functional effects of the mutation under different, controlled experimental conditions. Describing the dynamic effects caused directly by the mutation through a computational model yielded a novel mechanism of seizure susceptibility for this epilepsy patient, consistent with his phenotype.

Using these functional evaluations to achieve clinical improvements will require further study. Functional neurophysiological studies combined with computational modelling, illustrated in this case, show that rich information can be derived about abnormal neuronal dynamics for individual patients. Conducting these experiments on a larger scale will confirm whether mechanisms observed in individual patients are unique or will converge to common mechanisms which correlate to the phenotypes.

## Methods

### Clinical case report

This child was first admitted at the age of 6 months with a brief, self-terminating febrile seizure with a right-sided predominance of his twitching movements. He subsequently presented with prolonged recurrent seizures, both with and without fever, some lasting 30 minutes or more. These required emergency treatment with benzodiazepines, and one intensive care unit admission related to respiratory depression following treatment, as a consequence of which treatment with phenytoin was commenced, which reduced the duration of his seizures to less than 5 minutes. Interestingly, a proportion of these seizures were apparently provoked by a hot bath, or whilst playing in a very warm environment.

There was no evidence of focal neurological impairment after recovering fromseizures during any of his hospital admissions. He was born at term and had an uncomplicated perinatal course. There was no family history of epilepsy, neurodevelopmental or psychiatric conditions. No abnormalities were found on systemic examination and extensive cardiology review; his echocardiogram and electrocardiogram (ECG) were unremarkable.

Because of the clinical phenotype he underwent genetic sequencing of the *SCN1A* gene at 12 months of age. This showed a de novo heterozygous missense mutation (c.3818C>T) causing changes in a functionally significant and highly conserved region of the *SCN1A* protein (p.Ala1273Val). This genetic mutation, together with the clinical context suggests a diagnosis of a seizure disorder within the wider Dravet Syndrome spectrum.

Following his genetic diagnosis, his treatment was changed to Sodium Valproate, which he tolerates well and which has markedly reduced the number and duration of seizures. At the current time he continues to make age appropriate developmental progress.

### Bacterial Transformations and Mutagenesis

All bacterial transformations for this project were performed in TOP10/P3 *E. coli* bacteria (Invitrogen). As *SCN1A* is known to spontaneously mutate in bacterial cultures, the entire length of the sodium channel was sequenced following every transformation. Sequencing services were provided by Operon (Eurofins Genomics, AL, USA) and Genewiz (NJ, USA). The original *SCN1A* DNA in the PCDM8 vector was graciously provided by Dr. Lori Isom (University of Michigan). All transformants from the original *SCN1A* vector were found to express the T1967A and G5923T mutations (compared to NM_006920.4), which encode the V650E and A1969S mutants in NaV1.1, respectively. A QuikChange Lightning Mutagenesis kit (Agilent Technologies, CA, USA) with the following primers was used to get the WT and mutant DNA.

#### A1967T

5’ – CACTGTGGATTGCAATGGTG**T**GGTTTCCTTGGTTGGTGGAC - 3’

5’ – GTCCACCAACCAAGGAAACC**A**CACCATTGCAATCCACAGTG - 3’

#### T5923G

5’ – CTGATCTGACCATGTCCACT**G**CAGCTTGTCCACCTTCC - 3’

5’ – GGAAGGTGGACAAGCTG**C**AGTGGACATGGTCAGATCAG - 3’

#### C3818T

5’ – CTGGAAATGCTTCTAAAATGGGTGG**T**ATATGGCTATCAAAC - 3’

5’ – GTTTGATAGCCATAT**A**CCACCCATTTTAGAAGCATTTCCAG - 3’

### Electrophysiology

We grew CHOk1 cells (Sigma-Aldrich, MO, USA) in Ham’s F12 medium (Life Technologies, CA, USA) supplemented with 10% FBS (Life Technologies) at 37C in 5% CO2. 24-48 hours prior to experimentation we used Polyfect transfection reagent (Qiagen, Venlo, NL) to transfect cells with 1ug of the *SCN1A*, 1ug of eGFP, and 0.5ug of the β1 subunit using the protocols suggested by Qiagen. 8-12 hours after transfection, we plated cells on sterile glass coverslips.

Whole cell patch clamp experiments were performed at 32 °C, 37 °C, and 40 °C using borosilicate glass pipettes pulled with a P-1000 puller (Sutter Instruments, CA, USA), dipped in dental wax, and polished to a resistance of 1.0-1.5 MΩ. Extracellular solutions contained (in mM): 140 NaCl, 4 KCl, 2 CaCl2, 1 MgCl2, and 10 HEPES. Intracellular solutions contained (in mM): 130 CsF, 10 NaCl, 10 HEPES, and 10 EGTA. We titrated extracellular and intracellular solutions to pH 7.4 with CsOH.

We performed all experiments using an EPC9 patch-clamp amplifier digitized using an ITC-16 interface (HEKA Elektronik, Lambrecht, Germany). For data collection and analysis we used Patchmaster/Fitmaster (HEKA Elektronik) and Igor Pro (Wavemetrics, OR, USA) running on an iMac (Apple Inc., CA, USA). Temperature was maintained using a TC-10 temperature controller (Dagan Corporation, MN, USA). We low-pass-filtered the data at 5kHz and used a P/4 leak subtraction procedure for all recordings. The holding potential between protocols was −90mV.

### Pulse Protocols and Analysis

Macroscopic currents were elicited with 20ms depolarizations to membrane potentials between −100 mV and +60 mV. Conductance was determined by dividing peak current by the experimentally observed reversal potential subtracted from membrane potential. Normalized conductance plotted against voltage was fit by a single Boltzmann equation. The decay of current was fit by a single exponential equation to determine the time constant of open state inactivation at a given voltage.

Steady-state fast inactivation was measured as the proportion of current remaining in a test pulse to 0 mV following 200 ms pulses to voltages between −130 mV and +10 mV. The normalized current plotted against voltage was fit by a single Boltzmann equation.

The time course of fast inactivation recovery at −90 mV was measured as the proportion of current after a 200 ms depolarization to 0 mV and a recovery pulse of varying lengths to −90 mV. The normalized current was plotted versus recovery time and fit with a double exponential equation.

### Statistical Analysis

We used an analysis of covariance (ANCOVA) to test first for evidence that the A1273V mutant has differential temperature sensitivity compared to the WT channel. If there was no evidence for differences in temperature sensitivity, we tested whether the A1273V mutant differed from WT. In this analysis, the interaction between temperature (continuous variable) *vs*. mutant (nominal variable), was used as a predictor variable. A significant difference in the interaction term was evidence of a difference in temperature sensitivity between the mutant and WT channels. If this was not the case, then the analysis was repeated without the interaction term to test for a mutant effect. All statistical analyses were performed using JMP software (SAS Institute, NC, USA). Statistical significance was evaluated at P < 0.05. and measurements of error are reported as standard error of the mean.

### Modelling

Mutation effects on action potential generation were modelled using Hodgkin-Huxley (HH) models adapted to fit the dynamics of responses seen in regular spiking cortical pyramidal cells ^17^. The HH model estimates changes in membrane potential from non-linear, voltage-dependent changes in ion-specific membrane conductances:

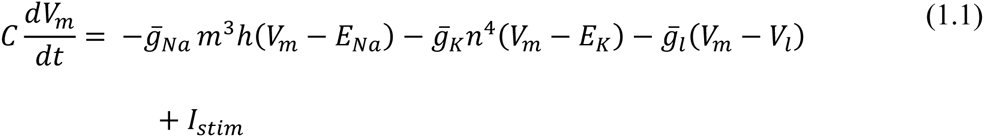

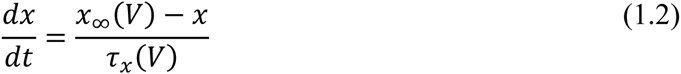

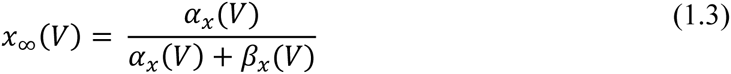

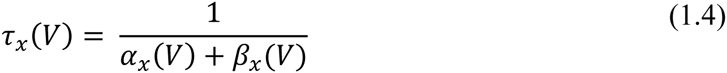

where *x* is *m*, *n*, or *h*

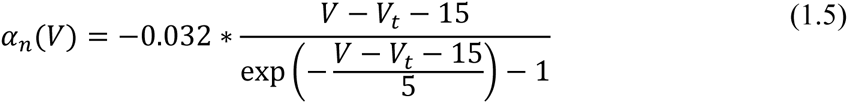

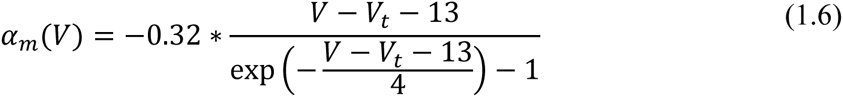

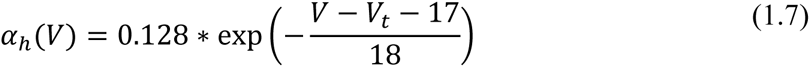

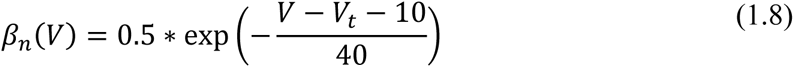

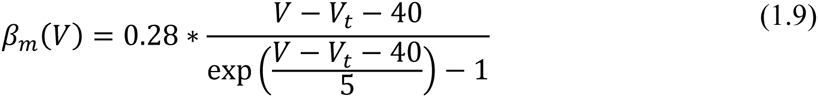

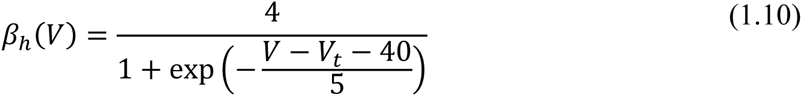

The equations represent a system of coupled ordinary differential equations describing the changes of membrane voltage (Eq. 1.1) and ion channel gating parameters (Eq 1.2). In this formulation there are only two ion channels — voltage gated-potassium channels, and voltage-gated sodium channels — the latter of which are of particular interest as we aim to parameterise a model that implements the experimentally derived changes in gating parameters *m* and *h* for the voltage gated sodium channel to estimate the resultant abnormal dynamics of the neuronal membrane.

At steady state, gating parameters are described by Eq. 1.3 ^26^. This represents a sigmoid function, which can also be parameterised using the generic Boltzmann formulation ^27^, Eq. 2.1:

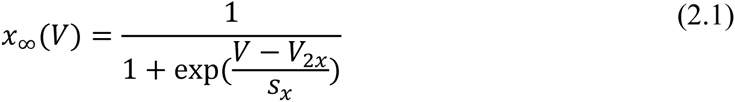

Experimental measurements taken from voltage clamp experiments include measures of midpoints (*V*_*2x*_) and slope (s_x_) of conductance and fast inactivation of sodium channels, representative of gating parameters *m* and *h* respectively, and thus speak naturally to the Boltzmann formulation of gating parameters. However, because the experimental design did not fully replicate physiological states in cortical neurons, we normalised our results to those described in ^17^ as follows:

1. Simulate cortical neurons using the parameterisation of ^17^.
2. Fit Boltzmann equations parameters to these simulations and derive midpoint voltages and slope of steady state gating parameters for the Pospischil parameterisation
3. For the WT simulations at 37°C, the parameters (V_2m_, s_m_, V_2h_, s_h_) were set to the same values as for the Pospischil parameterisation
4. For all remaining simulations, parameters were adjusted using the original parameterisation as baseline – preserving the absolute offset compared to the WT (V_2m_, V_2h_), or relative difference compared to the WT (s_m_, s_h_).
5. An additional offset parameter was introduced for fast inactivation gating equations 1.7 and 1.10 to account for temperature-dependent differences in time constants (note that with the Boltzman formulation for equation 1.3, only time constants (Eq 1.4) depend on equations 1.5 – 1.10).

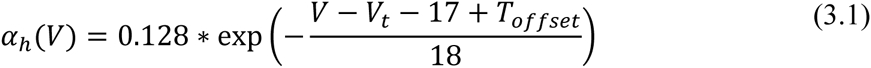

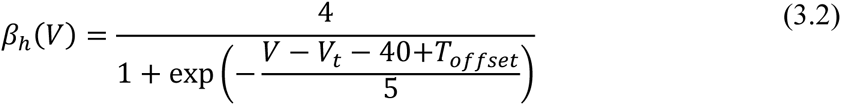

This yields four parameterisations of the same model, which represent different degrees of deviations from a standard model of regular spiking pyramidal cells. The normalisation was introduced to ensure that our experimental values are translated into physiological parameter spaces, but preserve the relative differences of the parameters between experimental conditions.

We examined the four versions of the model in terms of their response to different levels of stimulation (*I*_*stim*_, equation 1.1). This was first performed qualitatively and then further assessed using systematic variations and bifurcation analysis.

## Acknowledgements

We want to thank the patient and his family for agreeing to be part of this study. We also grateful to Deb K Pal for his helpful comments on the manuscript, as well as our funders: the Isaac Schapera Trust, the Wellcome Trust (RER), the Natural Sciences and Engineering Research Council of Canada, and the Canadian Foundation for Innovation.

